# Learning gene interactions from tabular gene expression data using Graph Neural Networks

**DOI:** 10.64898/2026.03.19.712949

**Authors:** Maria Boulougouri, Mohan Vamsi Nallapareddy, Pierre Vandergheynst

## Abstract

Gene interactions form complex networks underlying disease susceptibility and therapeutic response. While bulk transcriptomic datasets offer rich resources for studying these interactions, applying Graph Neural Networks (GNNs) to such data remains limited by a lack of methodological guidance, especially for constructing gene interaction graphs. We present REGEN (REconstruction of GEne Networks), a GNN-based framework that simultaneously learns latent gene interaction networks from bulk transcriptomic profiles and predicts patient vital status. Evaluated across seven cancer types in the TCGA cohort, REGEN outperforms baseline models in five datasets and provides robust network inference. By systematically comparing strategies for initializing gene–gene adjacency matrices, we derive practical guidelines for GNN application to bulk transcriptomics. Analysis of the learned kidney cancer gene-network reveals cancer-related pathways and biomarkers, validating the model’s biological relevance. Together, we establish a principled approach for applying GNNs to bulk transcriptomics, enabling improved phenotype prediction and meaningful gene network discovery.

## Introduction

Bulk RNA sequencing is one of the most widely used biological data types, where transcriptomic differences between conditions (e.g., healthy vs. disease) help identify perturbation-related genes. Traditional approaches, such as differential expression analysis, focus on individual genes, whereas functional and network-based methods aim to detect coordinated patterns across the transcriptome (1; 2). These methods assume that expression changes propagate through gene networks, allowing functionally related genes to be grouped into co-expression modules (3). Co-expression analyses, including widely used tools like WGCNA (4), have shown considerable success in identifying potential prognostic genes in multiple cancer types (5; 6).

Unlike traditional models that treat samples independently, graph neural networks (GNNs) leverage topological information. In gene expression data, connectivity often reflects interactions between gene products. GNN-based methods typically construct adjacency matrices from prior knowledge, including signaling pathways and protein–protein interaction (PPI) networks such as ConsensusPath DB (CPDB) (7), OmniPath (8), Human Protein Reference Database (9), Reactome (10), and most commonly STRING (11; 12). These networks, however, are not context-specific and may miss many gene regulatory relationships. Data-driven approaches, including matrix factorization (13) or topological analysis (14; 15), aim to learn informative graphs directly from the data. While some studies have compared co-expression and PPI-based input graphs (16), there is no clear consensus or comprehensive benchmark outlining how one chooses adjacency matrices for transcriptomic GNNs. Given recent work questioning GNN effectiveness for phenotype prediction (17), establishing clear guidelines is critical to efficiently apply these methods and reliably uncover latent gene relationships.

We hypothesize that learning the latent gene interaction graph can both improve model predictive power and uncover novel gene relationships. There are multiple methods to do so, but the three main classes of algorithms are: attention-based, graph learning–based, and dynamic graph–based. Attention-based methods learn edge and node-specific attention vectors to reconstruct the graph (18; 19; 20), but require an initial interaction matrix, which is often missing or sparse. Graph learning approaches design graphs as learnable parameters (21), via differentiable graph pooling (22; 23), or using graph kernel neural networks (24); however, these suffer from scalability issues. Dynamic graph methods, such as DGCNNs (25) and PGC-DGCNN (26), do not need prior connectivity knowledge as they rely on k-nearest neighbor (kNN) operations to build graphs.

Inspired by dynamic graph-based methods, we introduce REGEN (REconstruction of GEne Networks), which, to our knowledge, is the first graph inference framework to learn gene interactions from tabular bulk transcriptomic data. REGEN employs an efficient kNN-based method to perform graph-level classification tasks. We validate the method on synthetic data and conduct a thorough benchmark on seven real-world cancer datasets where it outperforms both structure-aware and structure-agnostic baselines. Through extensive ablation studies, we study how different adjacency initialization methods affect the model performance, and we provide guidelines for such applications. Finally, we conduct interpretability analyses on the REGEN learned graph for the kidney cancer dataset to reveal clusters and features enriched for cancer-related processes, including metabolic shifts, immunity, and known kidney cancer relevant genes.

## Results

In this study, we introduce a GNN-based framework, called REGEN, that classifies clinically relevant patient phenotypes, while inferring an optimal gene graph (fig. 1). We use bulk RNA sequencing from seven TCGA cancer datasets, where each patient is represented as a graph with genes as nodes and expression values as node features. Gene-to-gene adjacency matrices, including co-expression and prior knowledge -based information, can be optionally used as edge weights to further augment the graph. A GNN-based model is used to process these graphs for patient vital status classification. The model employs a kNN-based algorithm to learn an optimal graph of gene relationships through the training process, resulting in one final common gene connectivity network per cancer (see Methods).

**Figure 1.**
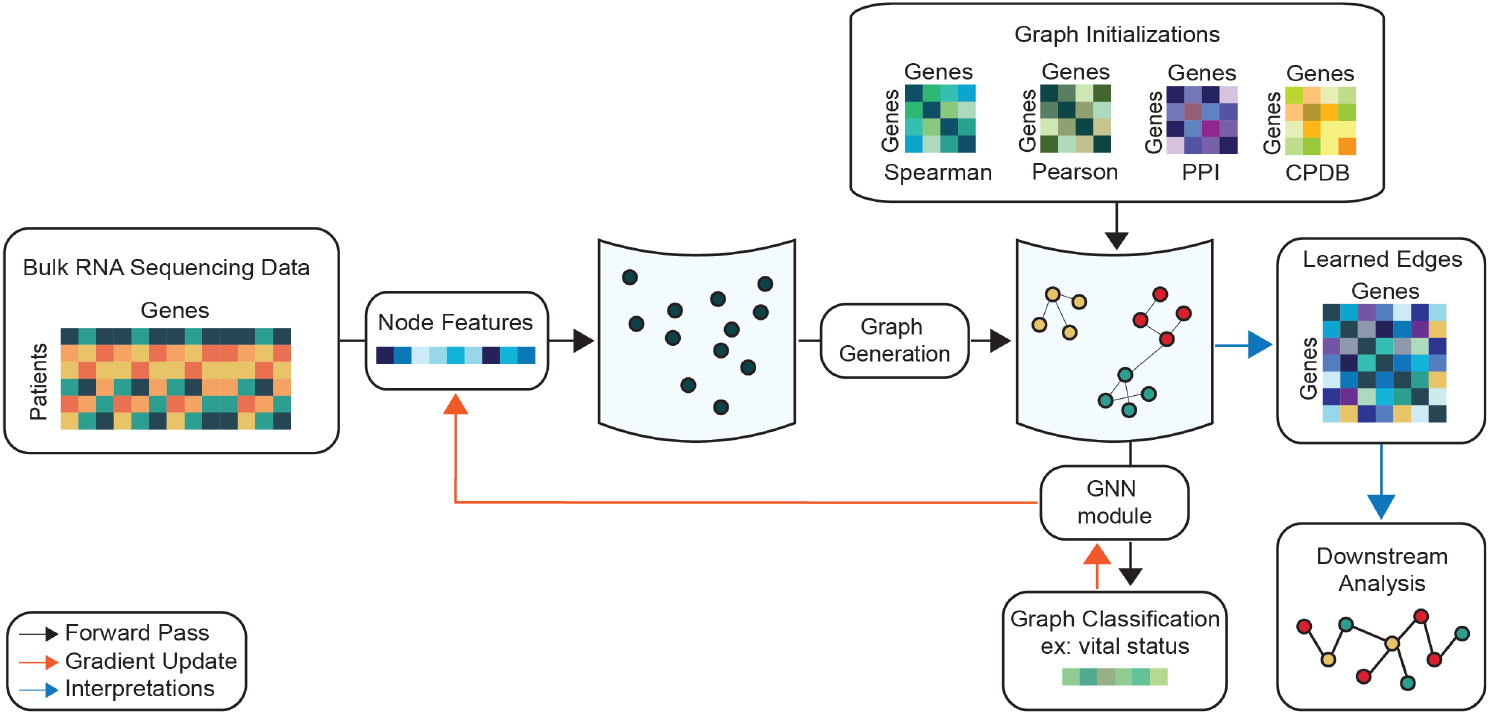
REGEN model overview. To generate the input graph, the genes are assigned as nodes, and the gene expression values are assigned as node features. The edges are generated by using a kNN assignment from the node embeddings. This graph is optionally augmented, in the form of edge weights, with information from pre-defined adjacency matrices, which could be based either on a transformation of the input data (Pearson, Spearman) or from prior interaction information (PPI, CPDB). Through subsequent iterations, the model updates the node embeddings, hence learning new edges. The model training is conducted end-to-end, and the new node features and edge connections are learned from the patient classification task. The final learned graph can be used for further downstream analysis to learn more about the interactions between different genes.

### REGEN effectively captures clusters in synthetic data

To validate whether REGEN can uncover latent node clusters, we first tested it on synthetic data where the ground truth labels are known. For this, we generated tabular data with sample-level binary labels following a shared graph structure. We generated two types of synthetic datasets with pre-defined clusters based on different types of graph connectivity: spatial or probabilistic (Exp 1 and 2, respectively) (fig. 2 (A)). Based on the sample label and cluster identity, node values were generated. The parameter signal strength (*ss*) was used to control the range of these values, and therefore the difficulty of the task (fig. 2 (B)). REGEN was run with default hyperparameters (table S1), and the output distance matrices were clustered and compared to ground truth (refer to Methods for further details on the experimental settings).

**Figure 2.**
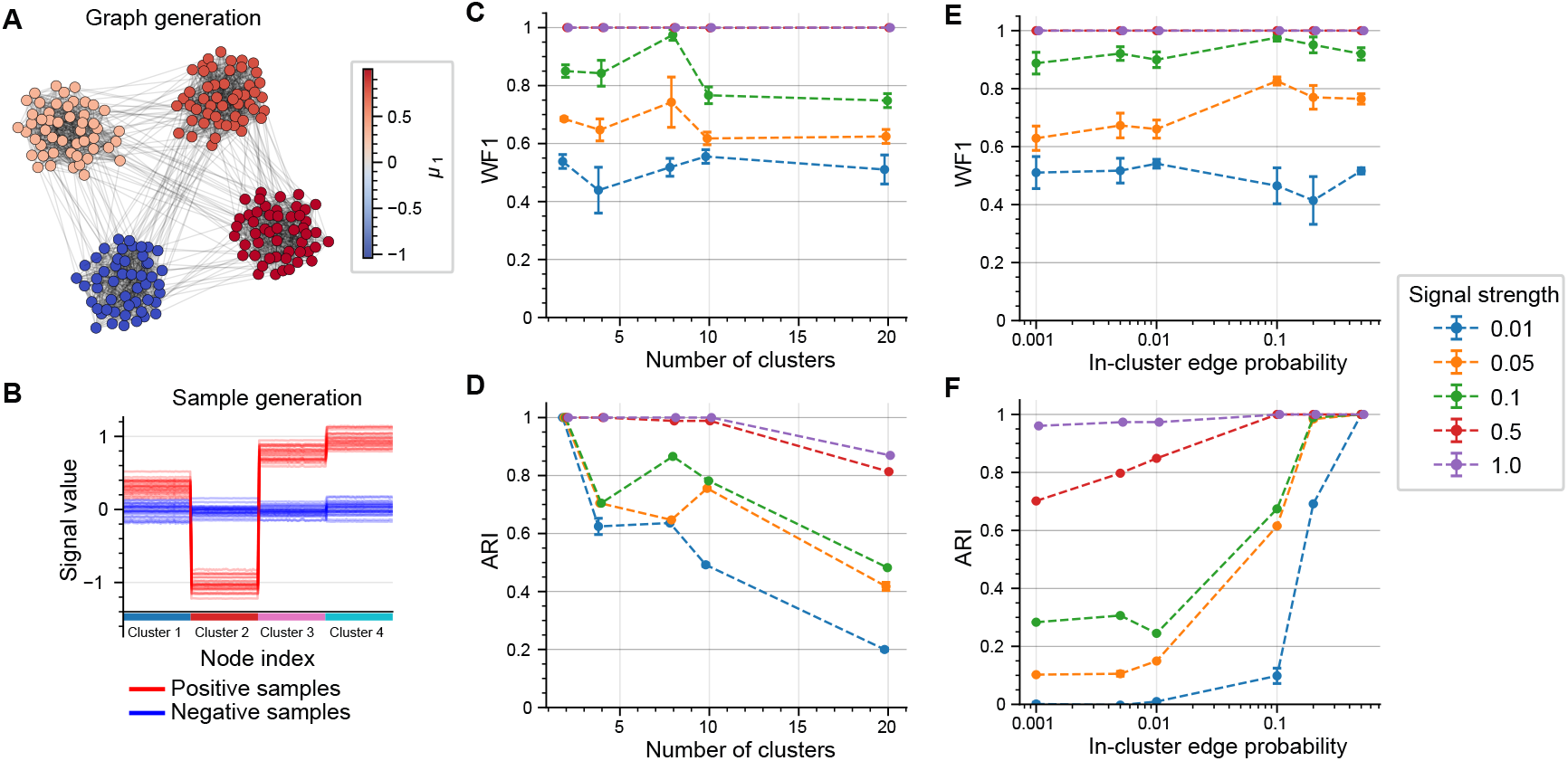
Overview of results from the synthetic data experiments. **(A)** Underlying graph generation, where nodes in different clusters have a different mean signal. **(B)** Individual node features are generated according to cluster and sample label. **(C)** Performance and **(D)** clustering quality results in Exp 1 with an increasing number of clusters. **(E)** Performance and **(F)** clustering quality in Exp 2 with increasing in-cluster edge probabilities compared to stable between-cluster edge probabilities.

In Exp 1, we notice that REGEN’s performance did not depend much on the cluster number, but rather on the *ss* parameter, which varied the WF1 scores between 50–100% (fig. 2 (C), table S2). Whereas the clustering performance depended on both the classification performance and the difficulty of the task. In the easiest task setting with 2 clusters, cluster assignments were perfect across all models, while in harder tasks with ≥ 10 clusters, only the highest performing models achieved higher clustering scores(fig. 2 (D), table S2). Consequently, we tested additional parameters on the 4 cluster setting with *ss* = 0.1. The signal smoothness is controlled by the heat parameter *t*; at a lower *t*, the signal was more localized, which resulted in a sharp drop in both classification and clustering, whereas a higher *t*, which indicates a smoother signal over the entire graph, improved classification without further enhancing clustering (table S3). Varying the sample labels by changing the positive label fraction (25%, 50%, 75%) had negligible effect (table S4). Interestingly, unequal cluster sizes improved both classification and clustering performance (table S5).

In Exp 2, the difficulty of each task was defined by the probabilities of edge occurrence between nodes within the same cluster (*p*_within_) and between different clusters (*p*_between_). With a stable *p*_between_ = 0.01 and 4 equally sized clusters, classification performance was again dependent on only signal strength (fig. 2 (E), table S6). Easier tasks, with high within-cluster connectivity (*p*_within_ = 0.5) led to perfect clustering regardless of signal strength (fig. 2 (F), table S6). Lower *p*_within_ resulted in clustering scores that followed model performance. When *p*_within_ ≤ *p*_between_, poorly performing models yielded random clustering, while even perfect classifiers achieved good but not perfect clustering.

Together, these results indicate that REGEN can accurately identify clusters from synthetic data. We note that not only can it detect clusters based on distances in the latent space, but it can also detect clusters based on graph connectivity.

### Benchmarking REGEN on real-world cancer data

To evaluate REGEN on real-world biological data, we used seven TCGA-derived cancer datasets: BRCA, KIPAN, GBMLGG, STES, HNSC, LUAD, and COADREAD (see Methods for data processing). We compared REGEN with both structure-aware models (GCNs) and a structure-agnostic baseline (MLP). Specifically, we implemented four GCN variants: GCN-Pearson, GCN-Spearman, GCN-PPI, and GCN-CPDB, based on the method used to construct the input adjacency matrix (see Methods for baseline architecture details). Across the seven cancer types, REGEN consistently outperforms the baseline models. It surpasses the MLP in five of the seven cancers and the GCN-based models in six of the seven (fig. 3 (A), refer to table S7 for WF1, table S8 for BalAcc). The performances of the REGEN models with the different types of information chosen for edge weight augmentations have been mentioned in table S9 (WF1), and table S10 (BalAcc). Additionally, when we average the normalized rankings across all cancers, we observe that REGEN achieves the best overall rank, followed by the MLP and then the GCN variants (fig. 3 (B)).

**Figure 3.**
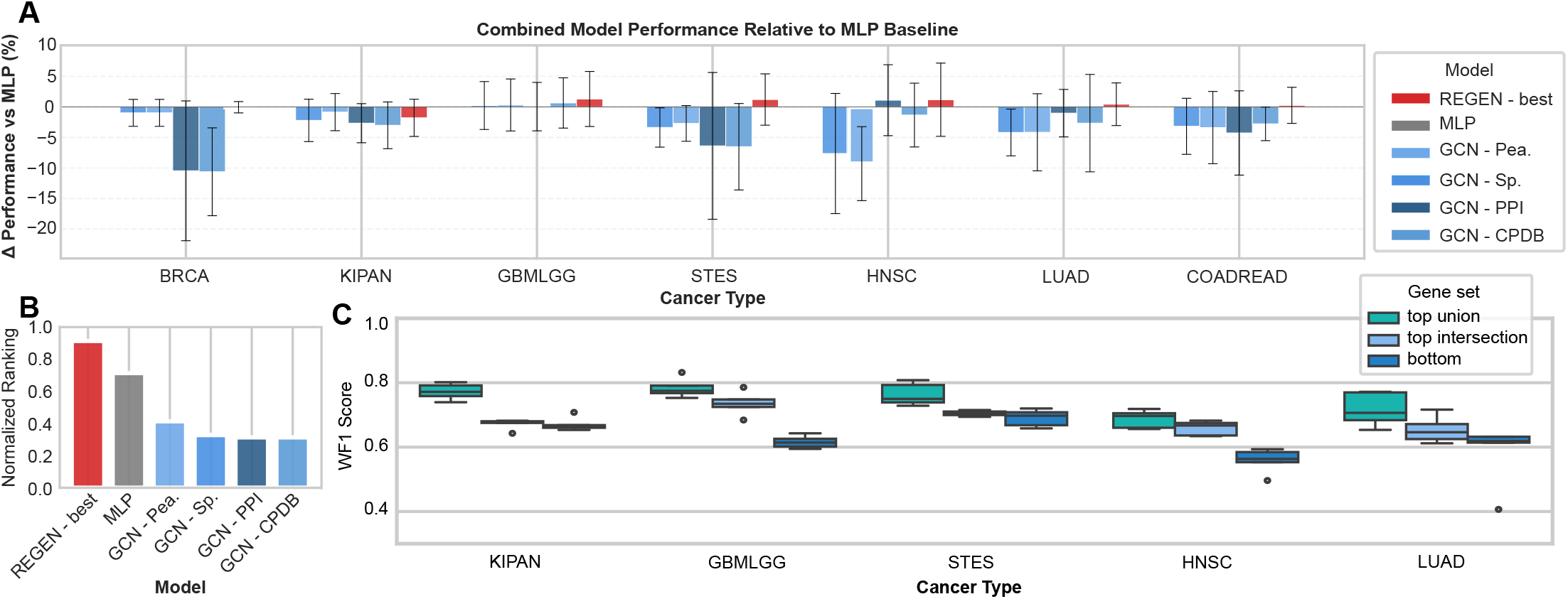
REGEN benchmarking and gene importance analysis. (**A)** Delta plot of model performances with MLP as a baseline. **(B)** Global normalized ranking plot comparing REGEN and the baselines. **(C)** WF1 scores of logistic regression models trained on three gene sets, top intersection, top union, and bottom, defined for the different cancers. The three gene sets were built using the node attributions obtained for all the genes in a dataset and choosing the intersection of the top 10% genes ranked by attribution per fold (top intersection), the union of these top 10% genes (top union), and a set of the bottom genes ranked by attribution of the same size as that of the top intersection gene set (bottom).

When comparing the performance of the GCN-based models in terms of WF1, we notice that prior knowledge–based graphs (PPI and CPDB) generally outperform correlation-based adjacency matrices (Pearson and Spearman) (tables S11 and S12). Similarly, we observe that REGEN benefits more from prior knowledge graphs as opposed to correlation-based information, achieving the highest performance in terms of WF1 for four of seven tasks (table S13). This together showcases that prior knowledge-based graphs extracted from PPI and CPDB generally provide more valuable information for the downstream tasks and can better help guide graph learning algorithms.

We observe that the best-performing GCN configurations are consistently outperformed by the MLP baseline, suggesting that static, predefined adjacency matrices may fail to capture the functional gene relationships needed for these prediction tasks. We observe that the choice of message-passing algorithm does not account for this gap as GCNs achieve the best classification performance in five of the seven tasks during REGEN’s hyperparameter search (table S13). This showcases a common challenge in applying GNNs to bulk transcriptomic data, which is that gene connectivity is often predefined rather than optimized for the task. By learning the adjacency structure directly from the data, we demonstrate that we can overcome this limitation and outperform both structure-aware and structure-agnostic baselines.

### REGEN identifies functional gene networks

To investigate what REGEN focuses on, we applied a standard gradient-based interpretability pipeline using the Integrated Gradients algorithm (27) to extract feature importance scores. These correspond to the relevance of individual genes to the prediction task (see Methods for details). We first analyzed gene importances by defining three gene sets: top union, top intersection, and bottom (see Methods for construction details). BRCA and COADREAD were excluded, as the former contained all genes in the top union and the latter had none in the top intersection. For the remaining five cancers, logistic regression models trained only using these gene sets showed that top union genes had the highest predictive performance, followed by top intersection and then bottom genes (fig. 3 (C)). These results indicate that REGEN effectively identifies genes most relevant for cancer-specific phenotype prediction.

We next examined the gene-gene connections in the learned REGEN graphs. Specifically, we looked at the graph learned by each of the REGEN models trained separately in the five training folds. The majority of the edges were present in all five folds across all seven datasets, indicating that the model tends to identify the same gene connections (fig. S1). For each of the seven datasets, we chose the edges that were present across all five REGEN folds and conducted a Fisher’s exact test comparing present versus absent edges against known protein-protein interactions. This test turned out to be significant for COADREAD, HNSC, and STES cancer datasets. A similar test using signaling pathway annotations from CPDB was significant for KIPAN and STES cancer datasets (table S14). These results highlight that REGEN is capable of reliably identifying edges that correspond to biologically meaningful gene relationships.

To evaluate whether the learned graph captures biologically meaningful clusters, we examined which genes were positioned close together in the embedding space. While a ground truth for gene-to-gene connectivity is unavailable, genes with shared functions, usually defined by gene ontologies, are expected to cluster together. As a proof of concept, we focused on the kidney cancer (KIPAN) dataset. We first extract gene embeddings from the best-performing fold of the REGEN model (Fold 4, WF1 = 82.20%, BA = 78.92%). The learned REGEN graph for this fold was clustered using the Leiden algorithm (resolution = 1.0, see table S15) and visualized using the UMAP algorithm (fig. 4 (A)). We then conducted an overrepresentation analysis on the obtained clusters to identify enriched pathways (see Methods for pipeline details). The three clusters that contained several general cancer and kidney cancer related pathways enriched were 15, 22, and 16 (table S16). Cluster 15 was found to be enriched for pathways with established roles in kidney cancer, including Hedgehog signaling (28), metabolism (29), and Epithelial-to-Mesenchymal Transition (EMT) (30), a process linked to fibrosis after kidney injury (31), malignant transformation of renal epithelial cells (32), Hedgehog signaling (33), and metabolic changes (34) (fig. 4 (B)). Cluster 22 was found with associations for adaptive and innate immunity, including antigen processing and presentation, NK cell-mediated cytotoxicity, and immunoregulatory interactions (fig. 4 (C)). Cluster 16 was identified to be enriched for pathways related to vesicle-mediated transport, autophagy, and organelle biogenesis and maintenance (fig. 4 (D)). Through this analysis, we demonstrate that REGEN is capable of organizing genes in the learned graph according to biologically meaningful functions, capturing important cancer-specific pathways.

**Figure 4.**
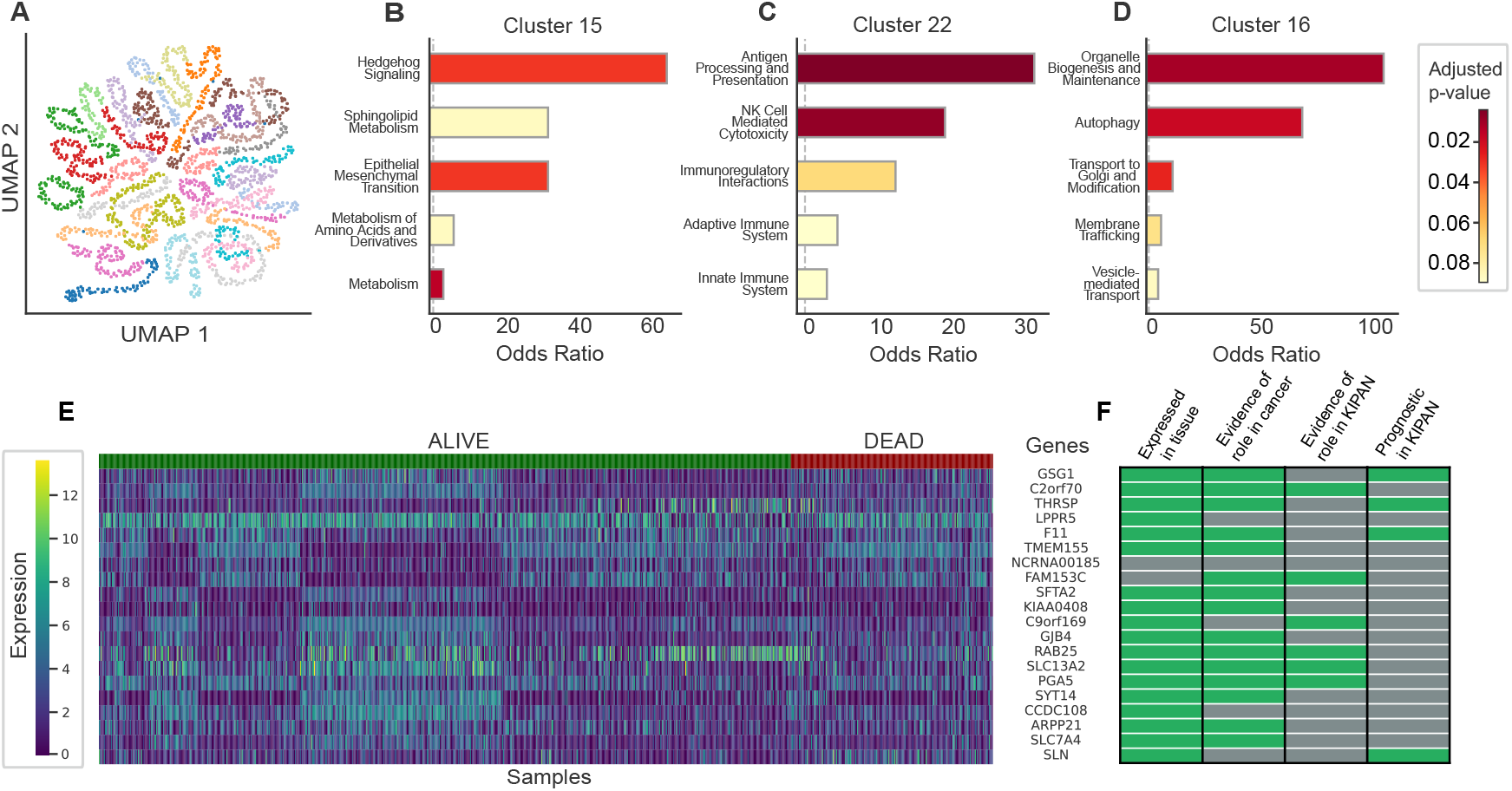
Biological interpretability analysis on the pan-kidney carcinoma (KIPAN) dataset. **(A)** UMAP visualization of the gene embeddings from the REGEN model, annotated with the cluster assignments from the Leiden algorithm. **(B-D)** Results of the overrepresentation analysis on selected clusters 15, 22, and 16. **(E)** Expression levels across patients of genes found in the top intersection gene set, and **(F)** evidence for these genes as possible biomarkers in terms of expression in kidney cancer tissue, evidence for gene involvement in cancer generally, or in KIPAN specifically, and evidence that the gene is prognostic for KIPAN.

To further assess whether REGEN captures biologically meaningful signals, we examined individual genes. For each training fold, we selected the top 10% of genes based on attributions and then focused on those that appeared in all five folds as potential biomarkers. For KIPAN, this resulted in a set of 20 genes. Their expression levels were visualized across alive and deceased patients (fig. 4 (E)). We evaluated these genes based on expression in kidney cancer tissue, evidence of involvement in cancer generally, specific roles in KIPAN, and prognostic value in KIPAN. Among them, *GSG1, THRSP, SLN*, and *F11* are previously identified prognostic markers of survival in kidney cancer patients (35), *C2orf70* and *FAM153C* are located under genomic regions significantly lost in kidney tumors (36), while *RAB25* (37), *SLC13A2* (38), *PGA5* (39), and *C9ORF169* (40) have evidence linking them to kidney cancer (fig. 4 (F)).

Overall, through these complementary analyses, we show that REGEN reliably identifies the most relevant genes and edges across multiple cancer types and, in the KIPAN case study, uncovers biologically meaningful structures and pathways, demonstrating its ability to capture both predictive and functionally relevant signals.

## Methods

### REGEN

In order to learn gene interaction relationships from different cancer-related datasets, we propose a graph-based machine learning model called REGEN, which stands for REconstruction of GEne Networks. The model consists of three components: (a) Embedding Module, (b) Graph Generation Module, and (c) Graph Classification Module (fig. 1). First, the input gene expression features are processed by the embedding module, which generates gene-level embeddings using a single linear layer. These embeddings are then passed to the graph generation module, which uses a kNN-based graph learning algorithm to construct a new adjacency matrix every forward pass. Genes with similar embeddings are connected via edges, forming clusters of genes with similar feature profiles. The resulting graph comprising the gene-level embeddings and kNN-derived edges is subsequently provided to the graph classification module. This module performs graph convolutions to predict downstream sample-level properties.

To guide the graph learning process, we assign edge weights based on gene connectivity derived from multiple sources. Specifically, we evaluate five configurations: (a) no weighting (None), (b) pairwise Spearman correlation across samples (Spearman), (c) pairwise Pearson correlation across samples (Pearson), (d) protein–protein interaction information from STRING (PPI) (41), and (e) shared signaling pathway membership from ConsensusPath DB (CPDB) (42). Hyperparameters of the REGEN model were optimized using Ray Tune (43), and the explored hyperparameter space is summarized in table S17. The model training was conducted using a stratified k-fold (*k* = 5) pipeline and a Binary Cross Entropy loss (see section S1.1). In order to compare the model performances, we used the Weighted F1 (WF1) and the Balanced Accuracy (BalAcc) metrics.

### Synthetic data

We generated two types of synthetic datasets based on an underlying graph connecting all features. In the first experiment (Exp 1), clusters were spatially separated. In the second experiment (Exp 2), clusters followed probabilistic connection patterns both within and between groups. For both experiments, the total graph consisted of 200 nodes. In Exp 1, we created random graphs with 2, 4, 8, 10, or 20 equally sized clusters. Nodes were sampled from Gaussian distributions centered at their cluster locations and connected based on spatial proximity, forming the ground truth binary adjacency matrix. In Exp 2, a stochastic block model (SBM) partitioned 200 nodes into clusters with connection probabilities *p*_within_ for nodes within a cluster and *p*_between_ for nodes between clusters.

From the adjacency matrix, we computed the graph Laplacian (*L*) and derived a covariance matrix via a heat kernel. Tabular data were generated with 1000 samples and randomly assigned binary labels. Node-level features were sampled from a Gaussian Random Field (GRF) defined by the covariance matrix, with the heat kernel formulation [*σ* = exp(−*tλ*)], where *t* is the diffusion parameter. A smaller *t* would produce localized correlations, while a larger *t* would diffuse them more broadly. For the positive class, mean feature values were shifted according to cluster structure, creating a learnable relationship between features and the graph. The mean depends on both cluster identity and class label, with the dependence strength controlled by the signal strength parameter *ss*, varied across [0.01, 0.05, 0.1, 0.5, 1] to simulate tasks of differing difficulty. To evaluate cluster assignment by REGEN, we extracted distance matrices for each training fold and applied agglomerative clustering (average linkage). Cluster similarity with the ground truth was measured using Adjusted Rand Index (ARI) and Normalized Mutual Information (NMI) metrics (see section S1.1).

### Data processing

We utilized datasets generated by the TCGA Research Network ^1^ with sufficient sample sizes (*>* 300 patients), which were obtained via the Firebrowse ^2^ portal (accessed 21.06.2024). Seven cancers were chosen from this portal, including breast invasive carcinoma (BRCA), colorectal adenocarcinoma (COADREAD), glioblastoma multiforme and lower-grade glioma (GBMLGG), head and neck squamous cell carcinoma (HNSC), pan-kidney carcinoma (KIPAN), lung adenocarcinoma (LUAD), and stomach and esophageal carcinoma (STES) cohorts. The clinical metadata, specifically patient vital status at the earliest follow-up (alive or dead), and RNA-Seq data from tumor sites were filtered for patients who have both types of information available. The downloaded Illumina HiSeq RNA-Seq expression data were normalized at the gene level and processed by the RSEM pipeline (44). Subsequently, genes with zero counts in more than 1% of the patients were filtered out, and a linear SVC with Lasso regularization (C=1000) was used to select the most relevant features. The expression values were *log*2(*x* + 1) transformed. The dataset statistics for all seven cancers can be found in table S18. The task is formulated at the patient level to predict the vital status after the first follow-up visit. This would be a binary classification task in nature.

### Benchmarks

We designed two types of baselines: structure-aware and structure-agnostic. The structure-agnostic baseline is a three-layer Multi-Layer Perceptron (MLP) with 64 nodes and 3 layers. For the structure-aware baseline, we used a two-layer Graph Convolutional Network (GCN). The GCN input adjacency matrix was static and binary in nature. This was constructed using four different approaches, resulting in four models: GCN-Pearson, GCN-Spearman, GCN-PPI, and GCN-CPDB. GCN hyperparameters were optimized using a grid search, with the search space detailed in table S19. For correlation-based methods, multiple correlation thresholds were evaluated. The correlation matrix was binarized by setting values within (mean ± std × thresh) to 0 and all others to 1. For prior-knowledge matrices, the presence of a protein–protein interaction (PPI) or co-membership of two genes in a CPDB pathway was encoded as 1, and absence as All benchmark models were trained using a pipeline similar to REGEN.

### Interpretability analysis

The proposed REGEN model uses gene embeddings to generate a kNN graph that is used as an adjacency matrix by the GNN-based classification module. To visualize the genes in the latent space, the gene embeddings were averaged across the sample dimension and extracted for the best-performing model fold. The resulting embeddings of the genes across the latent space dimensions were scaled and visualized on a 2D plot using UMAP (45) (n neighbors=20, min dist=5, metric = ‘euclidean’, spread=5).

To infer meaningful gene neighborhood information, the consensus graph consisting of the edges present across all 5 model folds was extracted and clustered using the Leiden algorithm implemented using the leiden package (46). The optimal *resolution parameter* was estimated by performing consensus clustering across resolutions ranging from 0.2 to 2.0. For each resolution, the Leiden algorithm was repeated 10 times independently with different random seeds. For each clustering partition, the modularity score was recorded, and the ARI was computed across all pairwise comparisons of clustering runs. The selected resolution balanced high modularity and stable clustering assignments (high mean ARI, low std ARI) across runs. Overrepresentation analysis (47) was performed on each cluster using the GSEApy package (48), with the total number of genes used for the training set as background. The genesets tested were GO Biological Process 2021 (49; 50), KEGG 2021 Human (51), Reactome Pathways 2024 (52), MSigDB Hallmark 2020 (53). Multiple testing correction according to the Benjamini-Hochberg method was performed on the p-values, and values below 0.1 were considered significant.

The REGEN model uses information from the learned gene features, in conjunction with the learned interactions between the genes, to conduct the classification task. To identify which genes are the most important for the classification task in the context of different cancers, a standard gradient-based interpretability technique called Integrated Gradients (27) was employed using Captum (54). Using this, we perturbed the learned graph to obtain node-level attributions for each of the samples in the datasets. For each training fold, the node attributions were extracted and ranked. The top 10% of genes are considered the most important per fold, with the genes appearing across all 5 folds being termed the “top intersection” genes and genes appearing in at least 1 of the 5 folds being termed the “top union” genes. We then rank-normalized the node attributions per fold; a set of genes of the same size as the “top intersection” genes with the lowest attributions overall were termed “bottom genes”.

## Conclusion

In this study, we introduced REGEN, a graph neural network–based framework designed for bulk transcriptomic data to infer gene–gene relationships. Across multiple cancer types, REGEN achieved performance comparable to or exceeding that of both structure-aware and structure-agnostic baselines while identifying biologically relevant genes. To address the relatively small size of typical transcriptomic datasets, REGEN employs a computationally efficient k-nearest neighbor (kNN)–based graph learning strategy. This design preserves model simplicity and interpretability while maintaining strong predictive performance. Our results further show that GCNs with static adjacency matrices often underperform structure-agnostic baselines, although their performance depends strongly on the choice of adjacency matrix and varies across datasets. In particular, prior knowledge–based matrices generally performed best across tasks, both when used as static graphs in GCNs and as edge weights in REGEN’s dynamic graphs. Overall, these findings suggest that tailored graph-based approaches such as REGEN can effectively capture biologically meaningful gene interaction patterns and provide valuable insights into disease mechanisms.

## Conflicts of interest

The authors declare that they have no competing interests.

## Data availability

The data underlying this article are available in the FireBrowse portal. GitHub for data analysis and model training: https://github.com/mariaboulougouri/REGEN.

## Author contributions statement

M.B. and M.V.N. conducted the experiments. M.B., M.V.N., and P.V wrote and reviewed the manuscript.

## Acknowledgments

This work is supported by funding from the Swiss National Science Foundation (SNSF: # 205884) and the grant for Graph Neural Networks for Explainable Artificial Intelligence ERA-NET + EJP (20CH21 195579).

## Footnotes

1 https://www.cancer.gov/tcga/

2 http://firebrowse.org/

## Notes

### Competing Interest Statement

The authors have declared no competing interest.

