## Supplemental Material for "Learning gene interactions from tabular gene expression data using Graph Neural Networks"

### Supplementary Material for manuscript: Learning gene interactions from tabular gene expression data using Graph Neural Networks

Maria Boulougouri, Mohan Vamsi Nallapareddy, Pierre Vandergheynst
LTS2 Signal Processing Laboratory, IEL-STI, EPFL, Rte Cantonale, 1015, Vaud, Switzerland

#### S1 Supplementary Methods

##### S1.1 Loss and Metrics

To measure the cluster similarity for the synthetic experiments, we use the Adjusted Rand Index (ARI) and the Normalized Mutual Information (NMI) scores. ARI corrects for chance, as defined in eq. (S1), where $RI$ is the Rand Index and $E[RI]$ its expectation for random clusterings. NMI measures similarity based on mutual information between clusterings $U$ and $V$ eq. (S2), with $H(U)$ and $H(V)$ denoting their respective entropies. The proposed REGEN model, GCN models, and the MLP were trained using the Binary Cross Entropy loss, defined in eq. (S3). For the evaluation of these models, we use the Weighted F1 score (WF1), and the Balanced Accuracy score (BalAcc).

$$\text{ARI} = \frac{RI - E[RI]}{\max(RI) - E[RI]} \tag{S1}$$

$$\text{NMI}(U, V) = \frac{I(U, V)}{\sqrt{H(U)H(V)}} \tag{S2}$$

$$\mathcal{L}_{\text{BCE}} = -\frac{1}{N}\sum_{i=1}^{N}\left[y_i \log(p_i) + (1 - y_i)\log(1 - p_i)\right] \tag{S3}$$

#### S2 Supplementary Figures

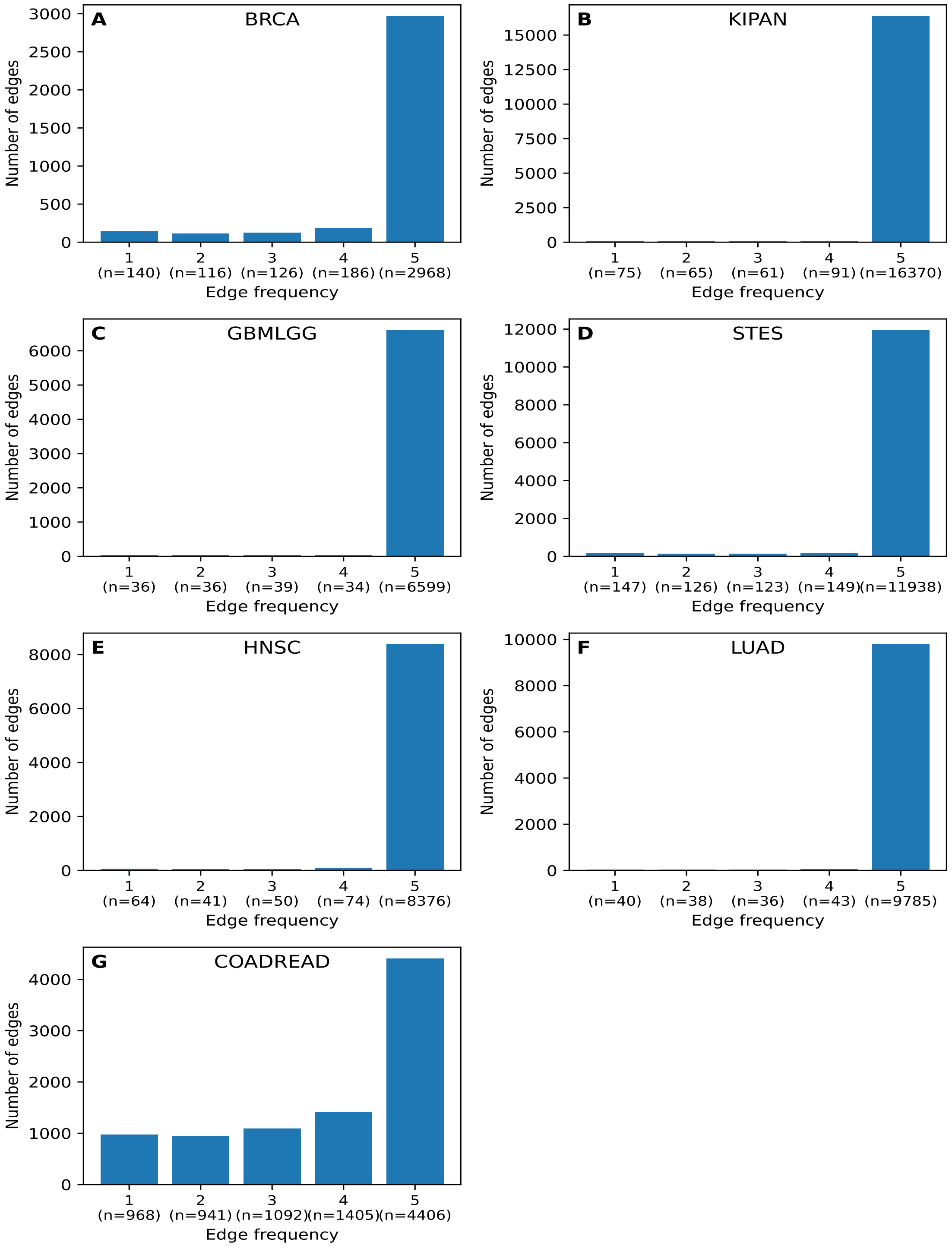


Figure S1: Edge consistency vs edge attribution across all cancer datasets. The number of REGEN training folds that an edge is found in is plotted on the x axis, with the total number of edges appearing in that number of folds found below. The averaged normalized edge attribution scores per edge are plotted on the y axis, with the majority of points around zero.

#### S3 Supplementary Tables

| Parameter | Value |
|---|---|
| **Initialization** | None |
| **GNN algorithm** | GCN |
| **Embedding size** | 64 |
| **No. of layers** | 2 |
| **No. of nodes** | 64 |
| **k neighbours** | 7 |
| **Pooling algorithm** | flatten |
| **Learning rate** | $1e-3$ |
| **Dropout value** | 0.2 |
| **Batch size** | 4 |
| **Distance metric** | cosine |

Table S1: Default hyperparameters set for REGEN to conduct the synthetic data experiments

| $n_{\text{clusters}}$ | Signal Strength | WF1 (%) | BA (%) | ARI | NMI |
|---|---|---|---|---|---|
| 2 | 0.01 | $53.83 \pm 2.39$ | $55.54 \pm 1.43$ | $1.0000 \pm 0.0000$ | $1.0000 \pm 0.0000$ |
| | 0.05 | $68.51 \pm 0.60$ | $68.63 \pm 0.68$ | $1.0000 \pm 0.0000$ | $1.0000 \pm 0.0000$ |
| | 0.1 | $85.01 \pm 2.15$ | $85.04 \pm 2.14$ | $1.0000 \pm 0.0000$ | $1.0000 \pm 0.0000$ |
| | 0.5 | $100.00 \pm 0.00$ | $100.00 \pm 0.00$ | $1.0000 \pm 0.0000$ | $1.0000 \pm 0.0000$ |
| | 1 | $100.00 \pm 0.00$ | $100.00 \pm 0.00$ | $1.0000 \pm 0.0000$ | $1.0000 \pm 0.0000$ |
| 4 | 0.01 | $43.92 \pm 7.91$ | $52.62 \pm 2.59$ | $0.6245 \pm 0.0280$ | $0.7718 \pm 0.0211$ |
| | 0.05 | $64.72 \pm 3.80$ | $65.17 \pm 3.75$ | $0.7047 \pm 0.0000$ | $0.8486 \pm 0.0000$ |
| | 0.1 | $84.26 \pm 4.46$ | $84.33 \pm 4.47$ | $0.7047 \pm 0.0000$ | $0.8486 \pm 0.0000$ |
| | 0.5 | $100.00 \pm 0.00$ | $100.00 \pm 0.00$ | $1.0000 \pm 0.0000$ | $1.0000 \pm 0.0000$ |
| | 1 | $100.00 \pm 0.00$ | $100.00 \pm 0.00$ | $1.0000 \pm 0.0000$ | $1.0000 \pm 0.0000$ |
| 8 | 0.01 | $51.81 \pm 3.09$ | $54.77 \pm 2.68$ | $0.6362 \pm 0.0000$ | $0.8076 \pm 0.0000$ |
| | 0.05 | $74.27 \pm 8.67$ | $74.62 \pm 8.42$ | $0.6470 \pm 0.0000$ | $0.8372 \pm 0.0000$ |
| | 0.1 | $97.50 \pm 1.87$ | $97.50 \pm 1.87$ | $0.8660 \pm 0.0000$ | $0.9524 \pm 0.0000$ |
| | 0.5 | $100.00 \pm 0.00$ | $100.00 \pm 0.00$ | $0.9884 \pm 0.0000$ | $0.9899 \pm 0.0000$ |
| | 1 | $100.00 \pm 0.00$ | $100.00 \pm 0.00$ | $1.0000 \pm 0.0000$ | $1.0000 \pm 0.0000$ |
| 10 | 0.01 | $55.53 \pm 2.35$ | $57.56 \pm 1.96$ | $0.4921 \pm 0.0055$ | $0.7127 \pm 0.0044$ |
| | 0.05 | $61.77 \pm 2.18$ | $63.09 \pm 1.59$ | $0.7558 \pm 0.0025$ | $0.8819 \pm 0.0015$ |
| | 0.1 | $76.66 \pm 2.88$ | $76.82 \pm 2.72$ | $0.7817 \pm 0.0000$ | $0.9063 \pm 0.0000$ |
| | 0.5 | $99.90 \pm 0.20$ | $99.90 \pm 0.20$ | $0.9887 \pm 0.0000$ | $0.9913 \pm 0.0000$ |
| | 1 | $100.00 \pm 0.00$ | $100.00 \pm 0.00$ | $1.0000 \pm 0.0000$ | $1.0000 \pm 0.0000$ |
| 20 | 0.01 | $51.05 \pm 5.00$ | $55.04 \pm 1.98$ | $0.1999 \pm 0.0046$ | $0.5519 \pm 0.0031$ |
| | 0.05 | $62.46 \pm 2.39$ | $63.34 \pm 2.44$ | $0.4176 \pm 0.0139$ | $0.7010 \pm 0.0072$ |
| | 0.1 | $74.83 \pm 2.40$ | $74.98 \pm 2.45$ | $0.4823 \pm 0.0000$ | $0.7601 \pm 0.0000$ |
| | 0.5 | $100.00 \pm 0.00$ | $100.00 \pm 0.00$ | $0.8134 \pm 0.0000$ | $0.9356 \pm 0.0000$ |
| | 1 | $100.00 \pm 0.00$ | $100.00 \pm 0.00$ | $0.8695 \pm 0.0000$ | $0.9604 \pm 0.0000$ |

Table S2: Results for the synthetic data "Exp 1", conducted with the parameters: $noiset = 1$, $n\_nodes = 200$, $n\_samples = 1000$, and $n\_labels = 2$ (equal distribution)

| Heat param | WF1 (%) | BA (%) | ARI | NMI |
|---|---|---|---|---|
| 0.01 | $64.52 \pm 1.17$ | $65.36 \pm 1.24$ | $0.2018 \pm 0.0000$ | $0.3385 \pm 0.0000$ |
| 0.1 | $72.55 \pm 2.21$ | $72.61 \pm 2.30$ | $0.6589 \pm 0.0000$ | $0.7658 \pm 0.0000$ |
| 0.5 | $86.89 \pm 4.19$ | $86.89 \pm 4.18$ | $0.6981 \pm 0.0000$ | $0.8302 \pm 0.0000$ |
| 1 | $84.26 \pm 4.46$ | $84.33 \pm 4.47$ | $0.7047 \pm 0.0000$ | $0.8486 \pm 0.0000$ |
| 2 | $88.60 \pm 2.31$ | $88.60 \pm 2.30$ | $0.7047 \pm 0.0000$ | $0.8486 \pm 0.0000$ |
| 5 | $98.90 \pm 0.73$ | $98.90 \pm 0.73$ | $0.7047 \pm 0.0000$ | $0.8486 \pm 0.0000$ |

Table S3: Results for the synthetic data "Exp 1" with varying values of the heat parameter ($t$). Here, the number of clusters was set to 4, and the signal strength ($ss$) to 0.1.

| **% of pos. label** | **WF1** | **BA** | **ARI** | **NMI** |
|---|---|---|---|---|
| 25 | 85.66 ± 2.89 | 83.69 ± 3.75 | 0.7047 ± 0.0000 | 0.8486 ± 0.0000 |
| 50 | 84.26 ± 4.46 | 84.33 ± 4.47 | 0.7047 ± 0.0000 | 0.8486 ± 0.0000 |
| 75 | 87.88 ± 3.51 | 87.88 ± 4.37 | 0.7047 ± 0.0000 | 0.8486 ± 0.0000 |

Table S4: Results for the synthetic data "Exp 1" with varying percentages of positive labels. Here, the number of clusters was set to 4, and the signal strength ($ss$) to 0.1.

| Cluster sizes | WF1 (%) | BA (%) | ARI | NMI |
|---|---|---|---|---|
| 50-50-50-50 | $84.26 \pm 4.46$ | $84.33 \pm 4.47$ | $0.7047 \pm 0.0000$ | $0.8486 \pm 0.0000$ |
| 90-50-50-10 | $99.40 \pm 0.20$ | $99.40 \pm 0.20$ | $0.9338 \pm 0.0000$ | $0.9292 \pm 0.0000$ |
| 10-90-10-90 | $90.89 \pm 2.07$ | $90.91 \pm 2.04$ | $0.8986 \pm 0.0000$ | $0.9001 \pm 0.0000$ |

Table S5: REGEN performance metrics across different cluster size distributions studied for synthetic data "Exp 1".

| $p_{\text{in}}$ | Signal Strength | WF1 (%) | BA (%) | ARI | NMI |
|---|---|---|---|---|---|
| 0.001 | 0.01 | $51.04 \pm 5.52$ | $54.42 \pm 1.13$ | $0.0012 \pm 0.0057$ | $0.0152 \pm 0.0050$ |
| | 0.05 | $62.88 \pm 4.19$ | $63.29 \pm 4.07$ | $0.1024 \pm 0.0000$ | $0.1665 \pm 0.0000$ |
| | 0.1 | $88.79 \pm 3.74$ | $88.82 \pm 3.71$ | $0.2833 \pm 0.0000$ | $0.3673 \pm 0.0000$ |
| | 0.5 | $100.00 \pm 0.00$ | $100.00 \pm 0.00$ | $0.7015 \pm 0.0000$ | $0.8331 \pm 0.0000$ |
| | 1 | $100.00 \pm 0.00$ | $100.00 \pm 0.00$ | $0.9606 \pm 0.0000$ | $0.9520 \pm 0.0000$ |
| 0.005 | 0.01 | $51.71 \pm 4.27$ | $54.19 \pm 1.33$ | $-0.0026 \pm 0.0002$ | $0.0143 \pm 0.0003$ |
| | 0.05 | $67.29 \pm 4.25$ | $67.42 \pm 4.09$ | $0.1055 \pm 0.0082$ | $0.1815 \pm 0.0020$ |
| | 0.1 | $92.10 \pm 2.35$ | $92.11 \pm 2.36$ | $0.3064 \pm 0.0000$ | $0.4117 \pm 0.0000$ |
| | 0.5 | $100.00 \pm 0.00$ | $100.00 \pm 0.00$ | $0.7973 \pm 0.0000$ | $0.8224 \pm 0.0000$ |
| | 1 | $100.00 \pm 0.00$ | $100.00 \pm 0.00$ | $0.9735 \pm 0.0000$ | $0.9646 \pm 0.0000$ |
| 0.01 | 0.01 | $47.67 \pm 5.09$ | $54.10 \pm 1.51$ | $0.0096 \pm 0.0013$ | $0.0324 \pm 0.0062$ |
| | 0.05 | $65.75 \pm 3.17$ | $66.05 \pm 3.13$ | $0.1495 \pm 0.0011$ | $0.2526 \pm 0.0039$ |
| | 0.1 | $89.97 \pm 2.68$ | $89.98 \pm 2.69$ | $0.2449 \pm 0.0000$ | $0.4275 \pm 0.0000$ |
| | 0.5 | $100.00 \pm 0.00$ | $100.00 \pm 0.00$ | $0.8486 \pm 0.0000$ | $0.8898 \pm 0.0000$ |
| | 1 | $100.00 \pm 0.00$ | $100.00 \pm 0.00$ | $0.9735 \pm 0.0000$ | $0.9696 \pm 0.0000$ |
| 0.1 | 0.01 | $46.49 \pm 6.20$ | $54.05 \pm 1.77$ | $0.0984 \pm 0.0264$ | $0.1390 \pm 0.0249$ |
| | 0.05 | $82.63 \pm 1.39$ | $82.75 \pm 1.26$ | $0.6152 \pm 0.0015$ | $0.7230 \pm 0.0066$ |
| | 0.1 | $97.60 \pm 1.20$ | $97.61 \pm 1.19$ | $0.6743 \pm 0.0000$ | $0.8031 \pm 0.0000$ |
| | 0.5 | $100.00 \pm 0.00$ | $100.00 \pm 0.00$ | $1.0000 \pm 0.0000$ | $1.0000 \pm 0.0000$ |
| | 1 | $100.00 \pm 0.00$ | $100.00 \pm 0.00$ | $1.0000 \pm 0.0000$ | $1.0000 \pm 0.0000$ |
| 0.2 | 0.01 | $41.43 \pm 8.23$ | $53.03 \pm 3.13$ | $0.6916 \pm 0.0000$ | $0.8119 \pm 0.0000$ |
| | 0.05 | $77.00 \pm 4.09$ | $77.22 \pm 3.96$ | $0.9840 \pm 0.0052$ | $0.9797 \pm 0.0051$ |
| | 0.1 | $95.10 \pm 2.73$ | $95.13 \pm 2.72$ | $0.9893 \pm 0.0054$ | $0.9858 \pm 0.0071$ |
| | 0.5 | $100.00 \pm 0.00$ | $100.00 \pm 0.00$ | $1.0000 \pm 0.0000$ | $1.0000 \pm 0.0000$ |
| | 1 | $100.00 \pm 0.00$ | $100.00 \pm 0.00$ | $1.0000 \pm 0.0000$ | $1.0000 \pm 0.0000$ |
| 0.5 | 0.01 | $38.43 \pm 3.93$ | $51.64 \pm 1.07$ | $1.0000 \pm 0.0000$ | $1.0000 \pm 0.0000$ |
| | 0.05 | $76.32 \pm 1.72$ | $76.45 \pm 1.77$ | $1.0000 \pm 0.0000$ | $1.0000 \pm 0.0000$ |
| | 0.1 | $91.99 \pm 2.11$ | $91.99 \pm 2.11$ | $1.0000 \pm 0.0000$ | $1.0000 \pm 0.0000$ |
| | 0.5 | $100.00 \pm 0.00$ | $100.00 \pm 0.00$ | $1.0000 \pm 0.0000$ | $1.0000 \pm 0.0000$ |
| | 1 | $100.00 \pm 0.00$ | $100.00 \pm 0.00$ | $1.0000 \pm 0.0000$ | $1.0000 \pm 0.0000$ |

Table S6: Results for the synthetic data ”Exp 2”, conducted with the parameters: $n\_clusters = 4$, $noiset = 1$, $n\_nodes = 200$, $n\_samples = 1000$, $p\_between = 0.01$, and $n\_labels = 2$ (equal distribution)

| **Model** | **BRCA** | **KIPAN** | **GBMLGG** | **STES** | **HNSC** | **LUAD** | **COADREAD** |
|---|---|---|---|---|---|---|---|
| **MLP** | **86.13 ± 0.88** | **80.05 ± 2.26** | 78.19 ± 2.84 | 74.00 ± 1.23 | 69.97 ± 4.73 | 72.73 ± 2.55 | 86.04 ± 1.92 |
| **GCN - Sp.** | 85.15 ± 2.02 | 77.84 ± 2.63 | 78.41 ± 2.70 | 70.63 ± 2.97 | 62.34 ± 8.61 | 68.54 ± 2.85 | 82.86 ± 4.16 |
| **GCN - Pea.** | 85.15 ± 2.02 | 79.19 ± 2.03 | 78.50 ± 3.17 | 71.29 ± 2.65 | 60.68 ± 3.77 | 68.57 ± 5.77 | 82.65 ± 5.59 |
| **GCN - PPI** | 75.67 ± 11.40 | 77.38 ± 2.26 | 78.23 ± 2.76 | 67.62 ± 11.94 | 71.06 ± 3.36 | 71.70 ± 2.93 | 81.77 ± 6.62 |
| **GCN - CPDB** | 75.51 ± 7.13 | 77.03 ± 3.08 | 78.83 ± 2.97 | 67.48 ± 6.96 | 68.63 ± 2.20 | 70.07 ± 7.55 | 83.26 ± 1.96 |
| **REGEN - best** | 86.07 ± 0.29 | 78.26 ± 2.04 | **79.48 ± 3.49** | **75.20 ± 4.01** | **71.14 ± 3.67** | **73.16 ± 2.38** | **86.29 ± 2.24** |

Table S7: Performance of the proposed REGEN model and the baselines across the seven different cancer datasets, measured in terms of the Weighted F1 Score (WF1). We notice that REGEN recovers performance lost in GNN-based models and achieves state-of-the-art performance in 5 out of 7 cancers. The best-performing model for each dataset has been mentioned in bold.

| **Model** | **BRCA** | **KIPAN** | **GBMLGG** | **STES** | **HNSC** | **LUAD** | **COADREAD** |
|---|---|---|---|---|---|---|---|
| **MLP** | 51.87 ± 2.51 | 68.42 ± 1.95 | 72.40 ± 3.44 | 58.60 ± 1.98 | 64.26 ± 5.32 | 59.12 ± 2.55 | 64.05 ± 2.96 |
| **GCN - Sp.** | 51.23 ± 1.69 | 71.45 ± 2.39 | 75.91 ± 2.58 | 56.24 ± 4.15 | 63.74 ± 3.44 | 53.56 ± 3.22 | 67.33 ± 9.14 |
| **GCN - Pea.** | 51.23 ± 1.69 | 72.19 ± 2.34 | 75.55 ± 2.42 | 55.94 ± 3.71 | 61.44 ± 2.85 | 62.13 ± 3.74 | 65.42 ± 7.94 |
| **GCN - PPI** | 64.52 ± 5.41 | 74.16 ± 1.96 | 76.12 ± 3.42 | 64.56 ± 2.77 | 68.52 ± 1.72 | 63.54 ± 4.56 | 73.38 ± 4.48 |
| **GCN - CPDB** | 61.11 ± 3.38 | 75.56 ± 1.64 | 76.16 ± 3.04 | 67.80 ± 3.33 | 65.65 ± 3.70 | 65.02 ± 6.74 | 73.63 ± 7.50 |
| **REGEN - best** | 50.38 ± 0.75 | 74.44 ± 2.61 | 77.91 ± 1.49 | 68.00 ± 2.04 | 69.60 ± 3.87 | 64.55 ± 5.14 | 74.38 ± 6.58 |

Table S8: Performance of the proposed REGEN model and the baselines across the seven different cancer datasets, measured in terms of the Balanced Accuracy (BalAcc).

| WF1 | BRCA | KIPAN | GBMLGG | STES | HNSC | LUAD | COADREAD |
|---|---|---|---|---|---|---|---|
| None | 85.92 ± 0.3 | 77.67 ± 2.25 | 78.16 ± 2.71 | 74.32 ± 5.44 | 66.95 ± 8.48 | 71.62 ± 2.49 | **86.29 ± 2.24** |
| Spearman | 84.98 ± 1.06 | 77.53 ± 3.42 | 78.49 ± 1.01 | 71.43 ± 2.08 | **71.14 ± 3.67** | 72.65 ± 2.83 | 85.85 ± 4.29 |
| Pearson | 81.27 ± 6.82 | 78.03 ± 3.71 | **79.48 ± 3.49** | 73.89 ± 3.46 | 68.35 ± 3.85 | 71.61 ± 3.60 | 84.37 ± 3.68 |
| PPI | **86.07 ± 0.29** | **78.26 ± 2.04** | 78.40 ± 3.63 | **75.20 ± 4.01** | 68.34 ± 2.70 | 71.50 ± 3.59 | 84.56 ± 6.61 |
| CPDB | 85.56 ± 0.93 | 78.04 ± 3.79 | 78.91 ± 3.66 | 74.80 ± 3.97 | 69.39 ± 6.62 | **73.16 ± 2.38** | 82.74 ± 5.72 |

Table S9: Performance of the best REGEN model in terms of WF1 depending on the choice of information used to supplement as edge weights. We notice that augmenting prior knowledge from PPI or CPDB as edge-weights results in higher performance in the majority of the datasets (4 out of 7).

| BA | BRCA | KIPAN | GBMLGG | STES | HNSC | LUAD | COADREAD |
|---|---|---|---|---|---|---|---|
| None | 50.22 ± 0.44 | 74.06 ± 2.77 | 75.86 ± 3.42 | 68.69 ± 5.25 | 67.61 ± 4.50 | 63.04 ± 5.36 | 74.38 ± 6.58 |
| Spearman | 53.98 ± 2.15 | 74.14 ± 2.45 | 76.38 ± 0.65 | 56.78 ± 2.35 | 69.60 ± 3.87 | 62.63 ± 4.98 | 75.46 ± 4.83 |
| Pearson | 56.08 ± 3.19 | 74.03 ± 2.45 | 77.91 ± 1.49 | 66.58 ± 2.83 | 66.02 ± 4.43 | 61.40 ± 2.32 | 73.69 ± 3.31 |
| PPI | 50.38 ± 0.75 | 74.44 ± 2.61 | 76.23 ± 4.35 | 68.00 ± 2.04 | 68.49 ± 2.26 | 60.65 ± 5.17 | 72.28 ± 3.73 |
| CPDB | 50.57 ± 1.13 | 75.66 ± 2.28 | 76.50 ± 2.26 | 69.85 ± 1.34 | 67.76 ± 2.16 | 64.55 ± 5.14 | 74.02 ± 6.99 |

Table S10: Performance of the best REGEN model in terms of BalAcc depending on the choice of information used to supplement as edge weights.

| Init | Threshold | BRCA | KIPAN | GBMLGG | STES | HNSC | LUAD | COADREAD |
|---|---|---|---|---|---|---|---|---|
| Spearman | 0.25 | 85.03 ± 2.25 | 74.18 ± 3.99 | 66.33 ± 4.88 | 69.77 ± 2.05 | 59.99 ± 5.50 | 59.73 ± 8.80 | 74.35 ± 9.56 |
| Spearman | 0.5 | **85.15 ± 2.02** | 75.86 ± 4.80 | 74.09 ± 1.09 | 69.77 ± 2.05 | 59.41 ± 3.95 | 57.74 ± 18.82 | 78.81 ± 5.81 |
| Spearman | 0.75 | 84.82 ± 2.65 | 76.36 ± 5.44 | 77.05 ± 3.96 | 69.65 ± 2.10 | 60.80 ± 4.12 | 68.54 ± 2.85 | 79.62 ± 11.42 |
| Spearman | 1.0 | 81.61 ± 4.41 | 76.92 ± 5.78 | 77.85 ± 4.12 | 70.63 ± 2.97 | 57.20 ± 2.65 | 57.66 ± 19.34 | 82.86 ± 4.16 |
| Spearman | 1.25 | 76.49 ± 16.92 | 77.84 ± 2.63 | 77.45 ± 4.06 | 63.62 ± 12.11 | 61.1 ± 5.22 | 59.01 ± 13.66 | 80.32 ± 5.11 |
| Spearman | 1.5 | 69.76 ± 16.73 | 77.01 ± 2.52 | 78.22 ± 4.24 | 65.08 ± 8.62 | 60.01 ± 6.05 | 61.75 ± 9.42 | 79.33 ± 10.54 |
| Spearman | 1.75 | 73.77 ± 7.04 | 77.65 ± 4.90 | 78.41 ± 2.70 | 66.78 ± 7.08 | 62.34 ± 8.61 | 68.05 ± 4.5 | 77.37 ± 6.86 |
| Pearson | 0.25 | **85.15 ± 2.02** | 73.39 ± 5.02 | 68.01 ± 3.87 | **71.29 ± 2.65** | 59.05 ± 5.66 | 60.19 ± 9.25 | 74.1 ± 9.92 |
| Pearson | 0.5 | 85.04 ± 1.96 | 75.36 ± 6.27 | 73.50 ± 3.22 | 70.01 ± 1.45 | 58.70 ± 5.29 | 56.69 ± 15.6 | 78.81 ± 5.81 |
| Pearson | 0.75 | 84.95 ± 2.41 | 76.73 ± 4.93 | 76.76 ± 4.48 | 68.21 ± 0.86 | 60.17 ± 5.57 | 62.11 ± 4.08 | 79.82 ± 11.52 |
| Pearson | 1.0 | 82.57 ± 4.48 | 78.63 ± 4.30 | 76.20 ± 4.11 | 66.37 ± 6.44 | 60.68 ± 3.77 | 67.1 ± 4.08 | 81.02 ± 3.71 |
| Pearson | 1.25 | 69.18 ± 16.87 | 78.53 ± 3.38 | 77.38 ± 2.78 | 64.53 ± 12.69 | 59.45 ± 8.07 | 55.59 ± 22.09 | 82.65 ± 5.59 |
| Pearson | 1.5 | 74.10 ± 12.46 | 77.48 ± 4.21 | 78.50 ± 3.17 | 67.56 ± 1.62 | 60.07 ± 5.34 | 63.07 ± 10.08 | 77.20 ± 5.36 |
| Pearson | 1.75 | 67.88 ± 13.18 | **79.19 ± 2.03** | 78.35 ± 3.35 | 69.47 ± 4.58 | 60.68 ± 5.7 | 68.57 ± 5.77 | 76.51 ± 7.71 |
| PPI | - | 75.67 ± 11.4 | 77.38 ± 2.26 | 78.23 ± 2.76 | 68.04 ± 4.45 | **71.06 ± 3.36** | **71.7 ± 2.93** | 81.77 ± 6.62 |
| CPDB | - | 75.51 ± 7.13 | 77.03 ± 3.08 | **78.83 ± 2.97** | 67.48 ± 6.96 | 68.63 ± 2.20 | 70.07 ± 7.55 | **83.26 ± 1.96** |

Table S11: Performance of the GCN-based models across the seven cancer datasets measured in terms of WF1. The Spearman and Pearson initialization matrices were binarized at different thresholds, and their performances have been reported separately. We notice that prior knowledge based GCNs using PPI or CPDB information for their adjacency matrix initializations result in a higher performance in the majority of the datasets (4 out of 7).

| Init | Threshold | BRCA | KIPAN | GBMLGG | STES | HNSC | LUAD | COADREAD |
|---|---|---|---|---|---|---|---|---|
| Spearman | 0.25 | 51.13 ± 1.51 | 66.85 ± 3.65 | 63.92 ± 1.4 | 53.84 ± 3.49 | 57.81 ± 3.72 | 52.52 ± 2.12 | 57.99 ± 4.31 |
| Spearman | 0.5 | 51.23 ± 1.69 | 66.91 ± 3.67 | 69.44 ± 4.66 | 53.84 ± 3.49 | 56.82 ± 1.9 | 54.77 ± 4.57 | 54.3 ± 1.98 |
| Spearman | 0.75 | 51.35 ± 1.91 | 69.35 ± 4.66 | 72.22 ± 5.51 | 53.73 ± 3.45 | 58.79 ± 3.69 | 53.56 ± 3.22 | 63.67 ± 8.34 |
| Spearman | 1.0 | 54.39 ± 3.76 | 70.77 ± 2.48 | 73.8 ± 4.52 | 56.24 ± 4.15 | 57.7 ± 5.09 | 55.42 ± 2.18 | 67.33 ± 9.14 |
| Spearman | 1.25 | 54.89 ± 3.63 | 71.45 ± 2.39 | 73.22 ± 6.36 | 58.46 ± 5.59 | 61.52 ± 2.55 | 51.32 ± 1.81 | 66.2 ± 10.04 |
| Spearman | 1.5 | 57.12 ± 5.8 | 71.78 ± 2.73 | 75.86 ± 4.23 | 62.44 ± 3.45 | 61.59 ± 2.98 | 59.57 ± 4.41 | 62.79 ± 8.23 |
| Spearman | 1.75 | 58.03 ± 3.71 | 71.66 ± 1.59 | 75.91 ± 2.58 | 63.19 ± 2.91 | 63.74 ± 3.44 | 60.42 ± 1.85 | 69.48 ± 6.4 |
| Pearson | 0.25 | 51.23 ± 1.69 | 66.91 ± 3.42 | 63.92 ± 1.4 | 55.94 ± 3.71 | 56.99 ± 4.65 | 58.08 ± 1.19 | 57.83 ± 4.07 |
| Pearson | 0.5 | 50.86 ± 1.72 | 67.43 ± 3.77 | 70.17 ± 4.73 | 54.71 ± 2.83 | 56.65 ± 4.57 | 56.78 ± 2.32 | 54.3 ± 1.98 |
| Pearson | 0.75 | 51.46 ± 2.1 | 70.3 ± 5.09 | 72.38 ± 5.57 | 51.68 ± 2.12 | 58.23 ± 4.1 | 52.58 ± 3.17 | 63.82 ± 8.46 |
| Pearson | 1.0 | 53.43 ± 3.88 | 73.92 ± 2.95 | 73.77 ± 4.9 | 58.79 ± 3.1 | 61.44 ± 2.85 | 54.97 ± 2.78 | 62.07 ± 6.31 |
| Pearson | 1.25 | 56.87 ± 2.55 | 72.75 ± 2.92 | 74.55 ± 3.6 | 58.28 ± 4.92 | 61.65 ± 2.52 | 51.44 ± 1.68 | 65.42 ± 7.94 |
| Pearson | 1.5 | 56.31 ± 4.63 | 73.3 ± 2.44 | 75.55 ± 2.42 | 59.7 ± 6.18 | 61.05 ± 3.38 | 59.93 ± 5.14 | 69.92 ± 5.77 |
| Pearson | 1.75 | 59.2 ± 3.57 | 72.19 ± 2.34 | 75.44 ± 3.46 | 63.79 ± 4.88 | 60.67 ± 2.09 | 62.13 ± 3.74 | 65.02 ± 7.57 |
| PPI | - | 64.52 ± 5.41 | 74.16 ± 1.96 | 76.12 ± 3.42 | 65.5 ± 2.69 | 68.52 ± 1.72 | 63.54 ± 4.56 | 73.38 ± 4.48 |
| CPDB | - | 61.11 ± 3.38 | 75.56 ± 1.64 | 76.16 ± 3.04 | 67.8 ± 3.33 | 65.65 ± 3.7 | 65.02 ± 6.74 | 73.63 ± 7.5 |

Table S12: Performance of the GCN-based models across the seven cancer datasets measured in terms of BalAcc. The Spearman and Pearson initialization matrices were binarized at different thresholds, and their performances have been reported separately.

| **Model** | **BRCA** | **KIPAN** | **GBMLGG** | **STES** | **HNSC** | **LUAD** | **COADREAD** |
|---|---|---|---|---|---|---|---|
| **Edge Weights** | PPI | PPI | Pearson | PPI | Spearman | CPDB | None |
| **GNN algorithm** | ChebNet | GCN | GCN | GCN | GCN | GCN | GAT |
| **No. of heads** | - | - | - | - | - | - | 8 |
| **Cheb filter** | 10 | - | - | - | - | - | - |
| **Embedding size** | 32 | 64 | 16 | 128 | 128 | 16 | 16 |
| **No. of layers** | 4 | 4 | 6 | 2 | 6 | 2 | 2 |
| **No. of nodes** | 32 | 128 | 128 | 32 | 128 | 32 | 64 |
| **k neighbours** | 3 | 13 | 7 | 9 | 7 | 9 | 3 |
| **Pooling algorithm** | topK | flatten | flatten | flatten | flatten | flatten | flatten |
| **Learning rate** | 0.00695 | 0.00094 | 4.9732e-05, | 0.00014 | 4.9732e-05 | 6.8513e-05 | 0.00122 |
| **Dropout value** | 0.17789 | 0.29190 | 0.09984 | 0.00547 | 0.09984 | 0.10928 | 0.26916 |
| **Batch size** | 2 | 4 | 1 | 8 | 1 | 2 | 32 |
| **Distance metric** | euclidean | cosine | cosine | euclidean | cosine | cosine | cosine |

Table S13: Best performing REGEN model configurations for the 7 cancer datasets obtained from the Ray Tune based hyperparameter optimization search. We notice that the majority of the time, the REGEN model prefers to be augmented with edge weights from prior knowledge extracted from PPI or CPDB. The GCN algorithm is most commonly chosen for optimizing performance. Additionally, the distance metric most commonly used to build the kNN graph was cosine.

| **Edges** | **BRCA** | **KIPAN** | **GBMLGG** | **STES** | **HNSC** | **LUAD** | **COADREAD** |
|---|---|---|---|---|---|---|---|
| **In graph and in PPI** | 106 | 505 | 230 | 405 | 285 | 297 | 122 |
| **In graph and not in PPI** | 2862 | 15865 | 6369 | 11533 | 8091 | 9488 | 4284 |
| **Not in graph and in PPI** | 49783 | 76571 | 40427 | 70393 | 67342 | 55723 | 140099 |
| **Not in graph and not in PPI** | 1544827 | 2514245 | 1321659 | 2806075 | 2157723 | 1805637 | 6547806 |
| **Fisher's exact test p-value** | 0.169 | 0.331 | 0.0149 | **2.47e-10** | **0.0476** | 0.789 | **0.00266** |
| **In graph and in CPDB** | 66 | 530 | 197 | 322 | 236 | 264 | 43 |
| **In graph and not in CPDB** | 2902 | 15840 | 6402 | 11616 | 8140 | 9521 | 1865 |
| **Not in graph and in CPDB** | 64746 | 77189 | 41799 | 64490 | 58957 | 48974 | 64769 |
| **Not in graph and not in CPDB** | 2820692 | 2513627 | 1320287 | 2811978 | 2166108 | 1812386 | 2821729 |
| **Fisher's exact test p-value** | 1 | **0.0556** | 0.721 | **0.00114** | 0.34 | 0.681 | 0.938 |

Table S14: Statistics on the number of PPI and CPDB edges that were present in the REGEN learned graph studied for all seven cancers.

| Resolution | Num. Clusters | | Modularity | | ARI | |
|---|---|---|---|---|---|---|
| | Mean | Std | Mean | Std | Mean | Std |
| 0.2 | 15.00 | 0.0000 | 0.9200 | 0.0006 | 0.9731 | 0.0252 |
| 0.4 | 20.60 | 0.4899 | 0.9330 | 0.0007 | 0.8965 | 0.0968 |
| 0.6 | 28.00 | 0.0000 | 0.9408 | 0.0001 | 0.9360 | 0.0685 |
| 0.8 | 31.80 | 0.4000 | 0.9427 | 0.0001 | 0.9846 | 0.0208 |
| 1.0 | 32.00 | 0.0000 | 0.9426 | 0.0000 | 0.9950 | 0.0101 |
| 1.2 | 34.00 | 0.0000 | 0.9422 | 0.0000 | 0.9925 | 0.0150 |
| 1.4 | 39.00 | 0.0000 | 0.9409 | 0.0000 | 0.9918 | 0.0088 |
| 1.6 | 41.00 | 0.0000 | 0.9399 | 0.0001 | 0.9787 | 0.0228 |
| 1.8 | 42.80 | 1.4697 | 0.9395 | 0.0009 | 0.9617 | 0.0358 |
| 2.0 | 45.00 | 0.0000 | 0.9382 | 0.0001 | 0.9640 | 0.0322 |

Table S15: Leiden clustering results across different resolutions conducted for the REGEN learned graph from the KIPAN dataset.

| Cluster ID | Gene set term | Gene set | Odds ratio | p value | Adjusted p value |
|---|---|---|---|---|---|
| Cluster 15 | Metabolism | Reactome Pathways 2024 | 3.687790906 | 0.01470635 | 0.01470635 |
| Cluster 15 | Epithelial Mesenchymal Transition | MSigDB Hallmark 2020 | 31.5 | 0.005668557 | 0.031177063 |
| Cluster 15 | Hedgehog Signaling | MSigDB Hallmark 2020 | 63.02857143 | 0.002893434 | 0.031177063 |
| Cluster 15 | Metabolism of Amino Acids and Derivatives | Reactome Pathways 2024 | 6.432352941 | 0.006029208 | 0.086820595 |
| Cluster 15 | Sphingolipid Metabolism | Reactome Pathways 2024 | 31.5 | 0.005668557 | 0.086820595 |
| Cluster 16 | Organelle Biogenesis and Maintenance | Reactome Pathways 2024 | 97.36764706 | 0.000113434 | 0.010946422 |
| Cluster 16 | Autophagy | Reactome Pathways 2024 | 63.97101449 | 0.002813901 | 0.018102763 |
| Cluster 16 | Transport to the Golgi and Subsequent Modification | Reactome Pathways 2024 | 10.77941176 | 0.005209686 | 0.027289127 |
| Cluster 16 | Membrane Trafficking | Reactome Pathways 2024 | 6.044117647 | 0.019626985 | 0.072846309 |
| Cluster 16 | Vesicle-mediated Transport | Reactome Pathways 2024 | 4.826470588 | 0.032829242 | 0.08679512 |
| Cluster 22 | Antigen processing and presentation | KEGG 2021 Human | 29.46666667 | 6.41954E-05 | 0.003980115 |
| Cluster 22 | Innate Immune System | Reactome Pathways 2024 | 3.715517241 | 0.009399516 | 0.089595978 |
| Cluster 22 | Adaptive Immune System | Reactome Pathways 2024 | 5.025287356 | 0.012559965 | 0.089595978 |
| Cluster 22 | Immunoregulatory Interactions Between a Lymphoid and a non-Lymphoid Cell | Reactome Pathways 2024 | 12.23888889 | 0.000799237 | 0.069488893 |
| Cluster 22 | Natural killer cell mediated cytotoxicity | KEGG 2021 Human | 18.39166667 | 0.00023662 | 0.00733523 |

Table S16: Results of over-representation analysis conducted on the REGEN learned graph from the KIPAN dataset.

| Parameter | Possible values |
|---|---|
| **Initialization** | None, Spearman, Pearson, PPI, CPDB |
| **GNN algorithm** | GCN, GAT, ChebNet, GINE, SAGE |
| **No. of heads** | 4, 8 |
| **Cheb filter** | 2, 4, 6, 8, 10 |
| **Embedding size** | 16, 32, 64, 128 |
| **No. of layers** | 2, 4, 6 |
| **No. of nodes** | 32, 64, 128 |
| **k neighbours** | 3, 5, 7, 9, 11, 13 |
| **Pooling algorithm** | mean, max, add, topk, asap, flatten |
| **Learning rate** | [1e-5, 1e-2] |
| **Dropout value** | [0.0, 0.3] |
| **Batch size** | 1, 2, 4, 8, 16, 32 |
| **Distance metric** | euclidean, cosine |

Table S17: Hyperparameter search space used for tuning the REGEN model.

| Statistic | BRCA | KIPAN | GBMLGG | STES | HNSC | LUAD | COADREAD |
|---|---|---|---|---|---|---|---|
| **No. of patients** | 1093 | 889 | 666 | 599 | 520 | 515 | 377 |
| **No. of genes** | 1788 | 2284 | 1655 | 2404 | 2114 | 1935 | 3659 |

Table S18: Statistics regarding the number of patient samples, and the genes for the seven TCGA cancer datasets after conducting the data processing pipeline.

| Parameter | Possible values |
|---|---|
| **Batch size** | $[2, 4, 8, 16]$ |
| **Num. Nodes** | $[32, 64]$ |
| **Learning Rate** | $[1e-2, 1e-3, 1e-4]$ |
| **Correlation threshold cut-off** | $[0.25, 0.5, 0.75, 1.0, 1.25, 1.5, 1.75]$ |

Table S19: Hyperparameter search space used for tuning the GCN-based models.
